## Supplemental Figures and Tables for "Noncanonical roles of chemokine regions in CCR9 activation revealed by structural modeling and mutational mapping"

### Supplementary Figures and Legends

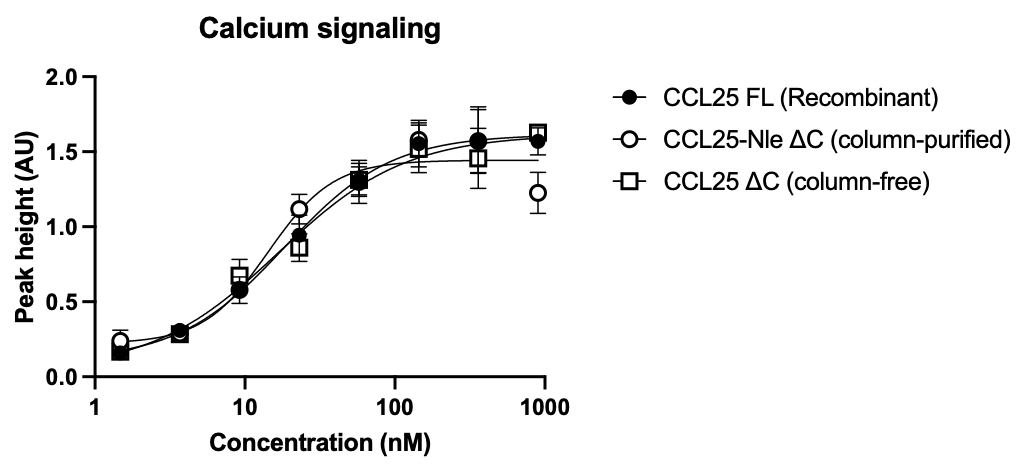

**Figure S1. Comparison between calcium flux response of MOLT-4 cells to full length (FL) recombinant CCL25 vs chemically synthesized C-terminally truncated CCL25 (ΔC) subjected or not to column purification (column-purified or column-free, respectively).** CCL25 variants were added to MOLT-4 cells at the indicated concentrations and Ca^2+^ flux signals were measured. Data points represent peak height (eq. 3) ± SEM from triplicate wells; the data shown are representative of 2 independent experiments.

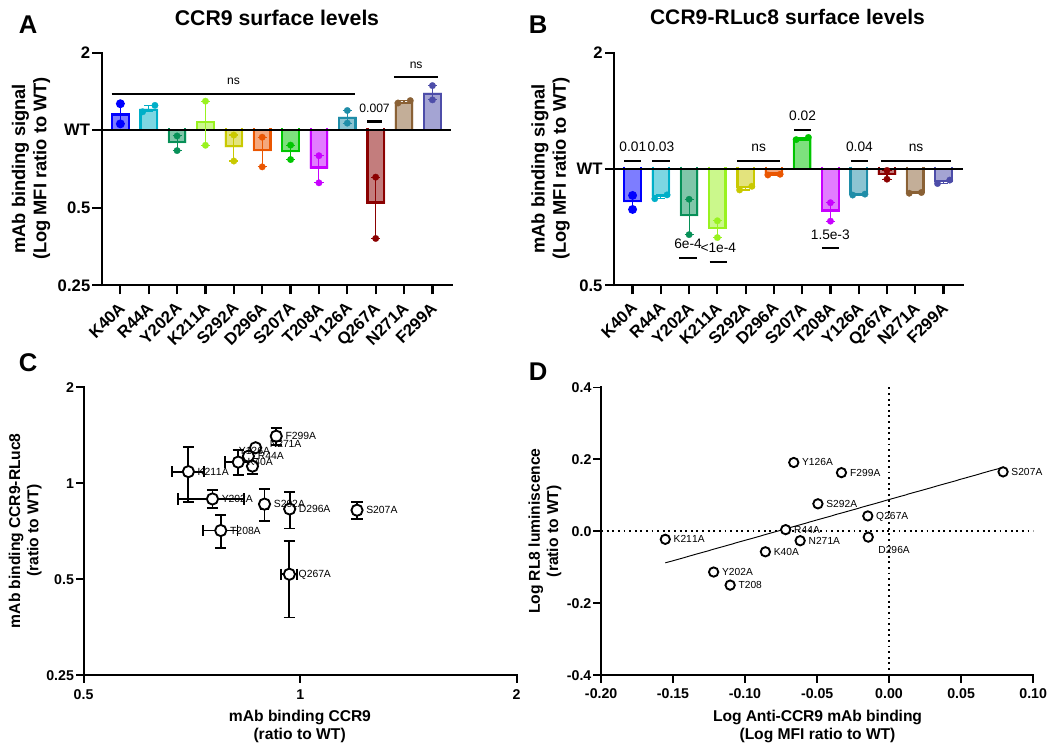

**Figure S2. Characterization of HEK-CCR9 and HEK-CCR9-RLuc8 YFP-Arr3 cell lines. A and B.** HEK293 cells expressing WT or mutant CCR9 (**A**) and HEK293 cells stably co-expressing YFP-Arr3 and WT or mutant CCR9-RLuc8 (**B**) were incubated with anti-CCR9 mAb at 4°C for 1h and analysed by flow cytometry. Signals are expressed as ratios to WT (eq. 1). Data represent mean ± SEM of 2 independent experiments and p-values were calculated using one-way ANOVA on log_2_ transformed ratios. **C.** Comparison between mAb binding to HEK-CCR9 untagged and HEK-CCR9-RLuc8 mutants shows no correlation indicating the lack of antibody sensitivity to the individual mutations. **D.** Comparison between luminescence and anti-CCR9 mAb binding, relative to WT receptor, for CCR9-RLuc8 mutants. Data points represent mean of luminescence (y axis) and mAb binding (x axis) relative to WT receptor of 2 independent experiments (simple linear regression: R^2^ = 0.3940, p-value = 0.0289).

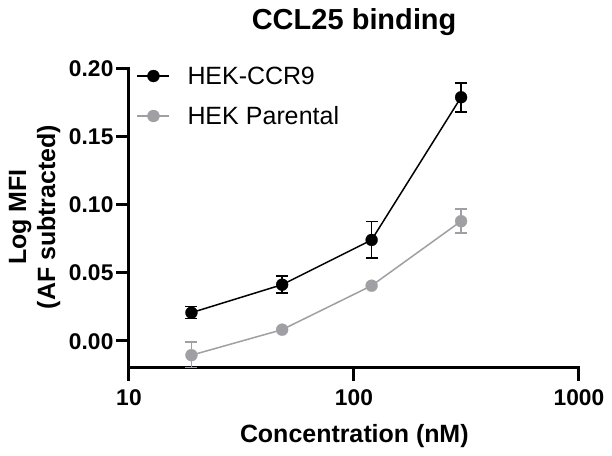

**Figure S3.** **Concentration-response curve for the binding of TAMRA-labelled CCL25 to HEK-CCR9 and HEK293 parental cell lines.** HEK293 parental cell line and HEK293 expressing WT CCR9 (HEK-CCR9) cells were incubated with TAMRA-labelled chemokine CCL25 at the indicated concentrations at 4°C for 1h. Binding signals are expressed as log MFI – log AF. Data represent mean ± SEM of triplicates in one experiment and are representative of 6 independent experiments.

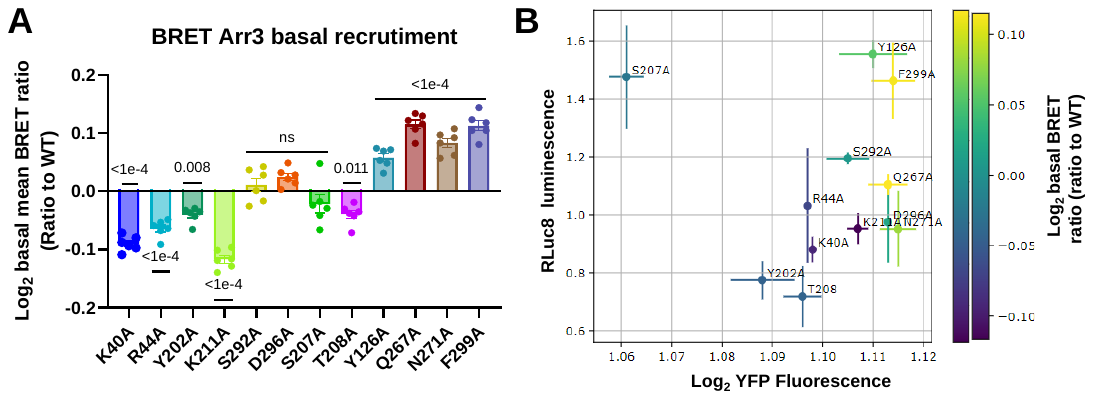

**Figure S4. Characterization HEK-CCR9-RLuc8 cell lines with respect to basal arrestin recruitment and acceptor:donor ratios. A.** HEK293 cells stably co-expressing YFP-Arr3 and WT or mutant CCR9-RLuc8 were incubated with BRET buffer containing 5 μM coelenterazine h and basal BRET signals were measured for 7 min. Data points represent the mean ± SEM of log_2_ transformed BRET signal normalized to WT receptor of 6 independent experiments. P-values were calculated using one-way ANOVA. **B.** Scatter plot of basal Arr3-YFP fluorescence vs basal CCR9-RLuc8 luminescence, both relative to WT cell line measured in the same experiment (log_2_ transformed axis). Plot points are colored according to basal BRET expressed as in panel **A**. Points represent mean ± SEM of RLuc8 luminescence (y axis) and YFP fluorescence (x axis) relative to WT receptor of 2 independent experiments.

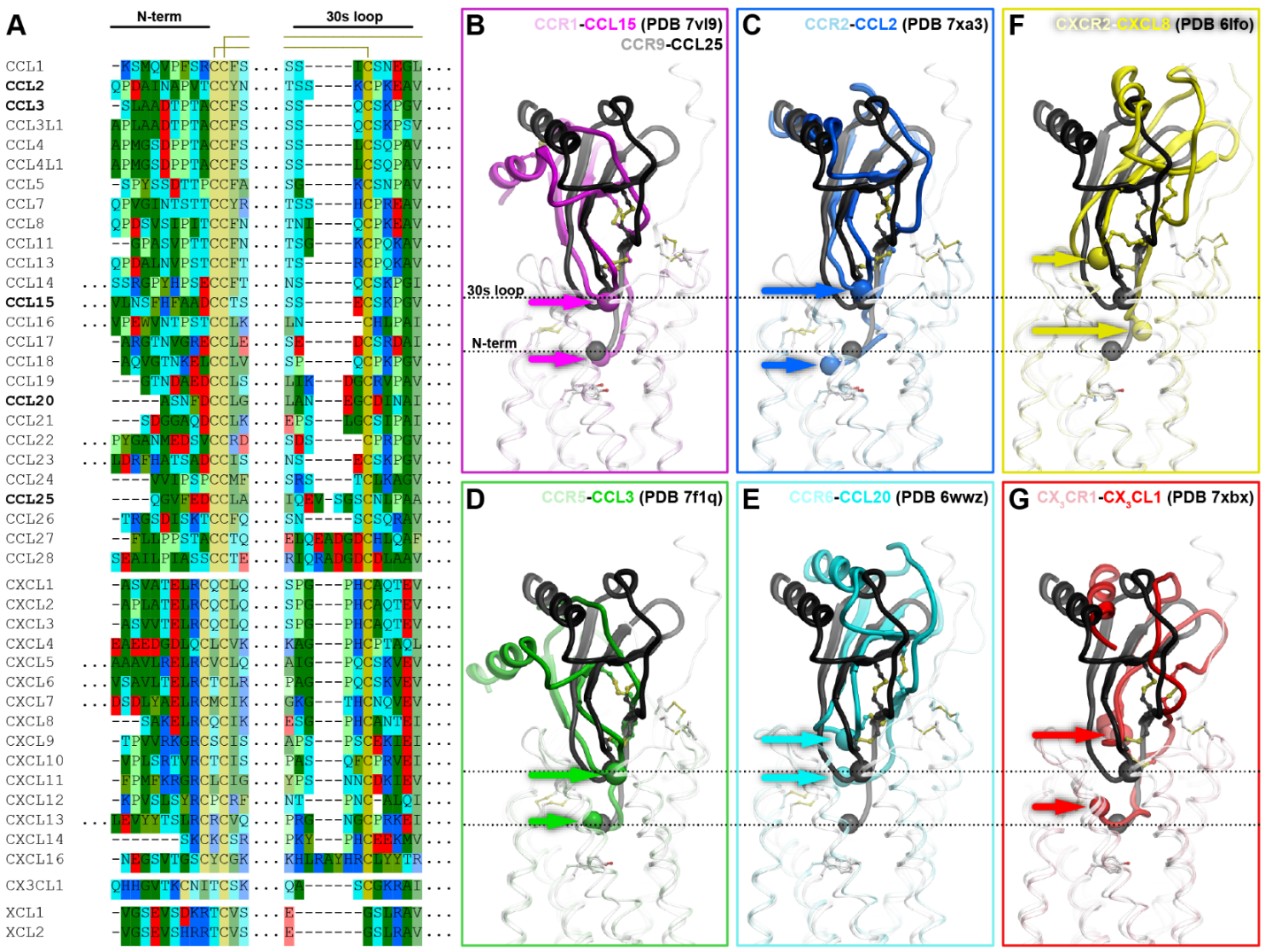

**Figure S5. Comparison of sequences and binding modes of CCL25 with other chemokines.** **A.** Partial sequence alignment of N-termini and 30s loops of human chemokines from the CC, CXC, CX3C, and XC subfamilies. Amino acid letters are colored by function (positive, negative, or neutral polar residues are highlighted in blue, red, and cyan respectively; non-polar residues in green, proline in light green, and cysteine in yellow). Chemokines whose experimental structures are shown in B-G are bold. **B-G**. Superimpositions of the predicted CCR9-CCL25 complex (grey and black ribbons) with selected experimental structures of other chemokine-receptor complexes (colored ribbons). Complexes are superimposed by receptor TM domains to depict variation in binding depths of chemokine N-termini and 30s loops (colored arrows) in reference to those of CCL25 (dashed lines). For each chemokine structure shown, C-alpha positions of the N-terminal residue and the residue preceding the third (30s loop) cysteine are denoted with spheres.

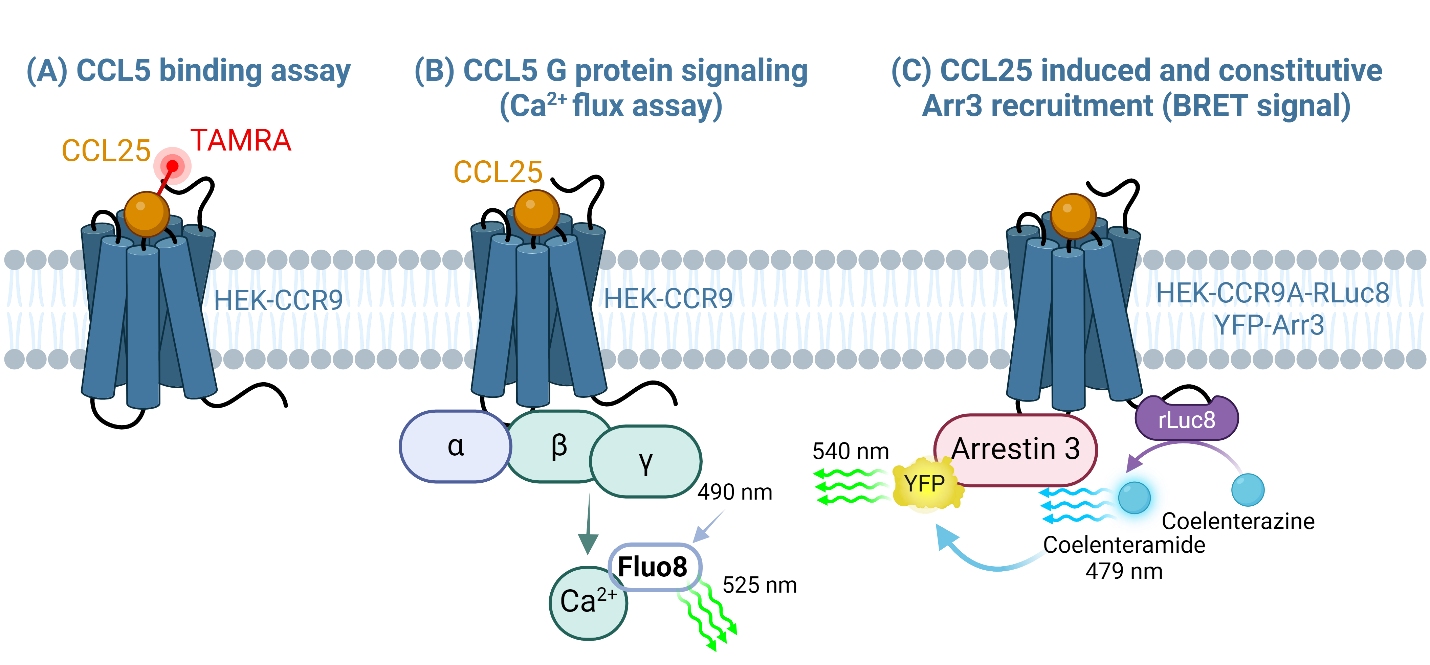

**Figure S6. Strategy for evaluating the roles of CCR9 residues in chemokine binding and signaling.** Evaluation of the impact that the 12 selected CCR9 CRS2 mutations have on CCR9 function, including their ability to bind CCL25 (**A**), to promote intracellular calcium (Ca2+) mobilization (**B**), and to recruit arrestin 3 in response to CCL25 (**C**).

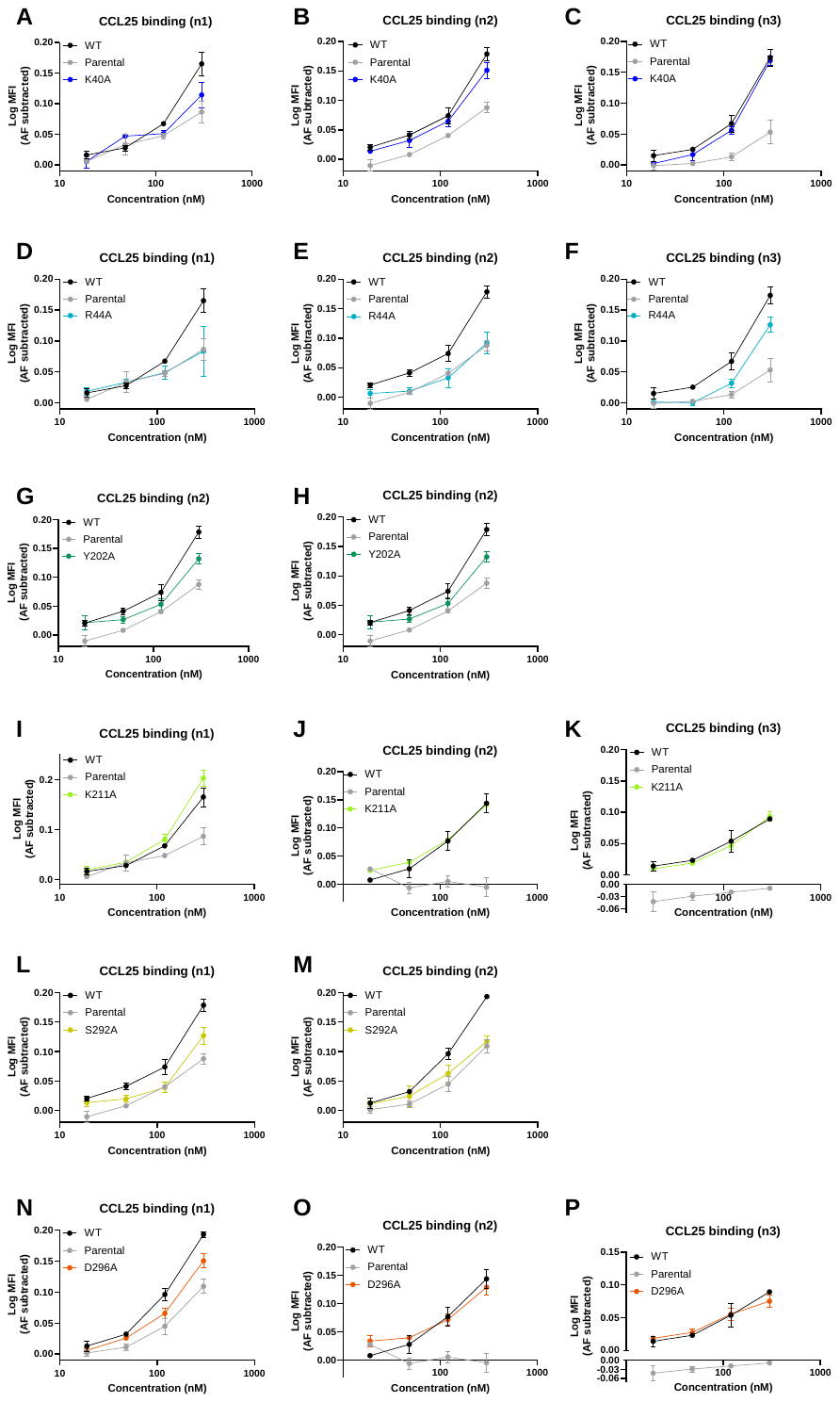

**Figure S7. Concentration-response curves for CCL25 binding to K40^1.24^A (A-C), R44^1.28^A (D-F), Y202^ECL2^A (G-H), K211^5.35^A (I-K), S292^7.28^A (L-M) and D296^7.32^A (N-P) CCR9 mutants compared to CCR9 WT.** HEK293 parental cell line (parental) and HEK293 cells expressing WT or mutant CCR9 cells were incubated at 4°C for 1h with TAMRA-labelled chemokine CCL25 at serial dilutions (300, 120, 48, 19 nM). Binding signals are expressed as log MFI – log AF where AF (autofluorescence) is the MFI of the corresponding cell line in the same experiment in the absence of the fluorescent chemokine. Data points represent mean ± SEM of triplicates in one experiment. 2 to 3 independent experiments are shown for each mutant.

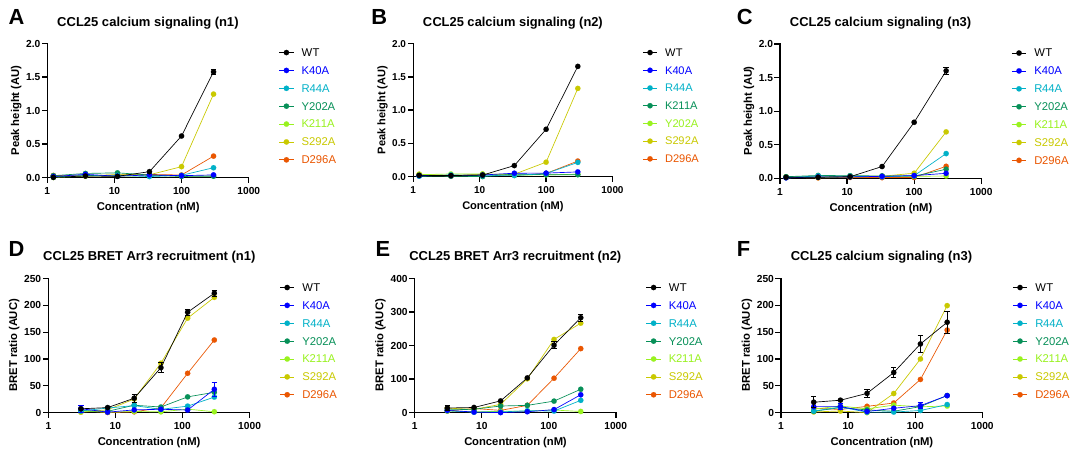

**Figure S8. Concentration-response curves for CCL25-induced Ca^2+^ flux (A-C) and BRET Arr3 recruitment (D-F) in K40^1.24^A, R44^1.28^A, Y202^ECL2^A, and K211^5.35^A, S292^7.28^A and D296^7.32^A CCR9 mutants compared to CCR9 WT. A-C.** CCL25-induced Ca^2+^ signaling on WT CCR9 or CCR9 mutants expressed in HEK293 cells. Ca^2+^ signals in response to CCL25 at the indicated concentrations are shown as mean peak height (**eq. 3**) ± SEM from triplicate wells. **D-F.** BRET assays for CCL25-induced Arr3 recruitment on WT CCR9 and CCR9 mutants. Data points represent mean ± SEM of normalized BRET signal AUCs (**eq. 4**) obtained in triplicate wells. **A-F.** 3 independent experiments are shown per mutant.

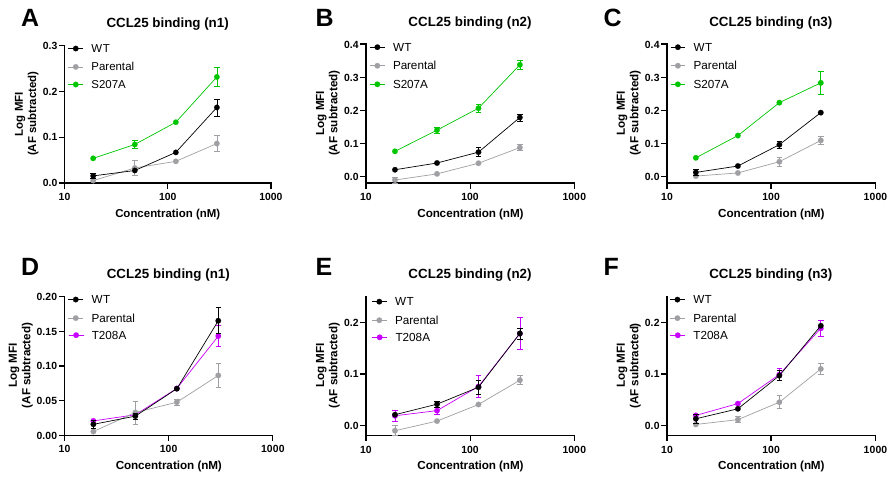

**Figure S9. Concentration-response curves for CCL25 binding on S207^5.31^A (A-C) and T208^5.32^A (D-F) CCR9 mutants compared to CCR9 WT.** Refer to **Fig. S7** legend for details.

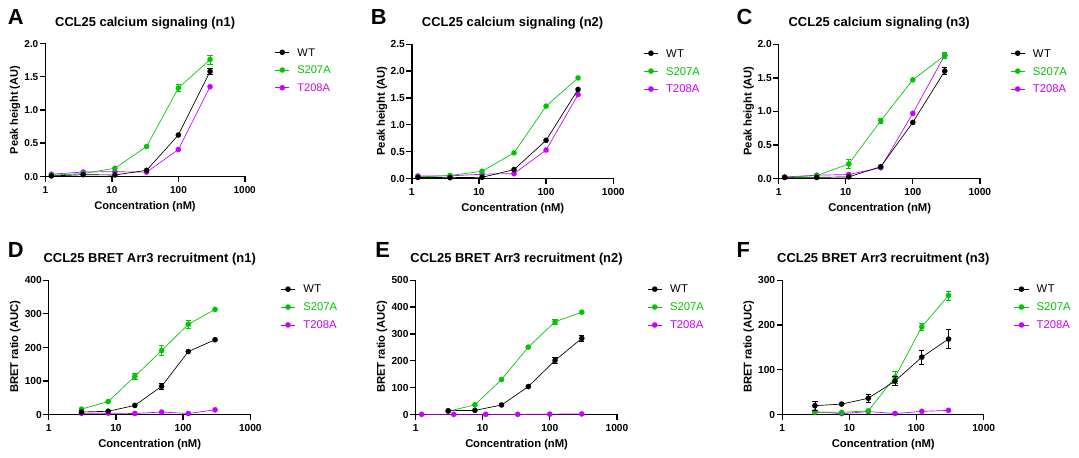

**Figure S10. Concentration-response curves for CCL25-induced Ca^2+^ flux signaling (A-C) and BRET Arr3 recruitment (D-F) on S207^5.31^A and T208^5.32^A CCR9 mutants compared to CCR9 WT.** Refer to **Fig. S8** legend for details**.**

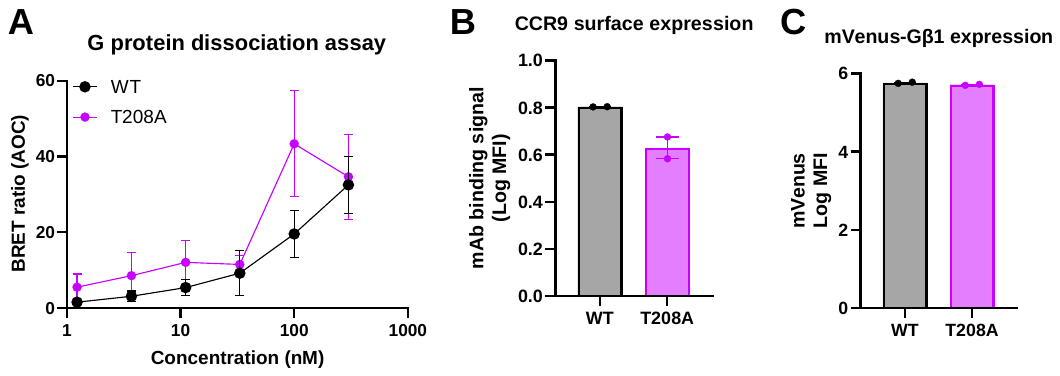

**Figure S11. G protein dissociation BRET assay on HEK-CCR9-T208^5.32^A. A.** HEK293 cells expressing WT or mutant CCR9 (T208^5.32^A) co-transfected with Gαi(91)-Rluc2, mVenus-Gβ1 and Gγ2 were stimulated with CCL25 at the indicated concentrations and BRET signals were measured. Data points represent mean ± SEM of area over the curve (AOC) of the BRET (eq. 4) obtained in technical triplicates in one experiment and are representative of 2 independent experiments. **B.** HEK293 cells expressing WT or mutant CCR9 (T208^5.32^A) were incubated with anti-CCR9 mAb at 4°C for 1h and analysed by flow cytometry. Signals are expressed as log*MFI*_WT or T208A_ – log*MFI*_parental_. Data represent mean log MFI of duplicates ± SEM. **C.** Transfection efficiency of mVenus-Gβ1 was evaluated by measuring mVenus fluorescence by flow cytometry. Signals are expressed as log*MFI*_WT or T208A_ – log*MFI*_parental_. Data represent mean log MFI of duplicates ± SEM.

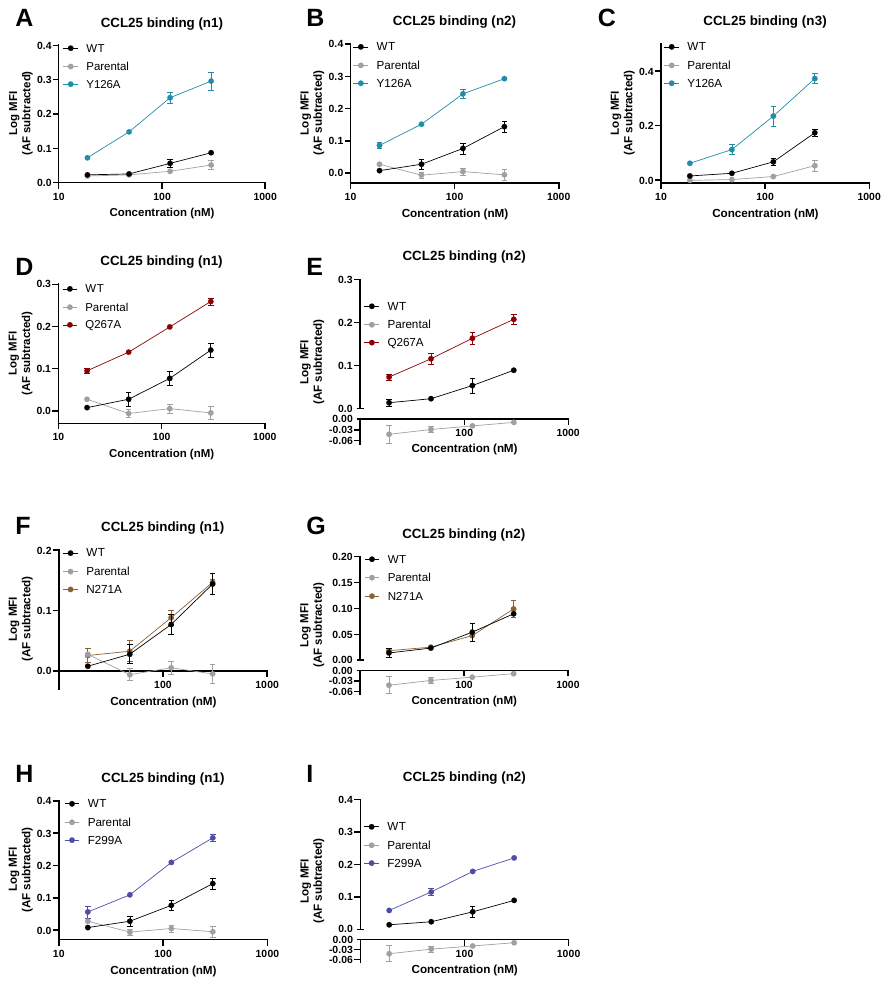

**Figure S12. Concentration-response curves for CCL25 binding to Y126^3.32^A (A-C), Q267^6.48^A (D-F), N271^6.52^A (G-I), and F299^7.35^A (J-L) CCR9 mutants compared to CCR9 WT.** Refer to **Fig. S7** legend for details.

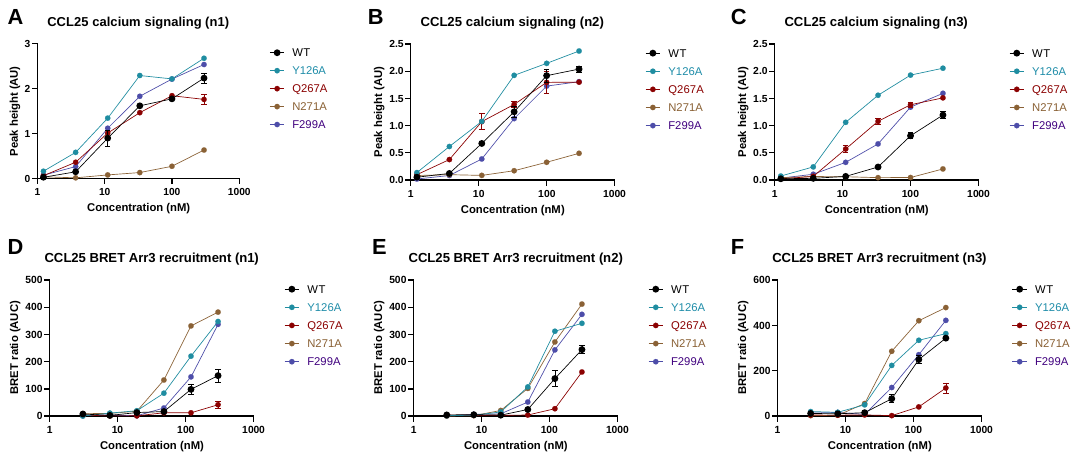

**Figure S13. Concentration-response curves for CCL25-induced Ca^2+^ flux signaling (A-C) and BRET Arr3 recruitment (D-F) on Y126^3.32^A, Q267^6.48^A, N271^6.52^A, and F299^7.35^A compared to CCR9 WT. A-C.** Refer to **Fig. S8** legend for details.

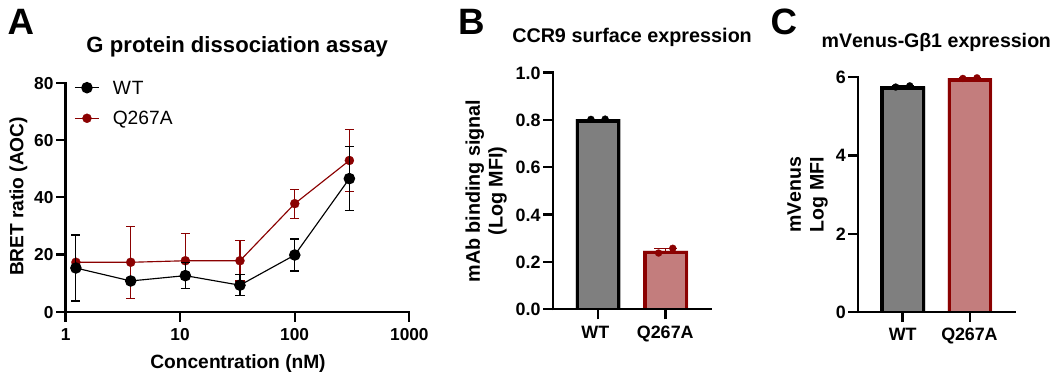

**Figure S14. G protein dissociation BRET assay on HEK-CCR9-Q267^6.48^A.** Refer to **Fig. S11** legend for details.

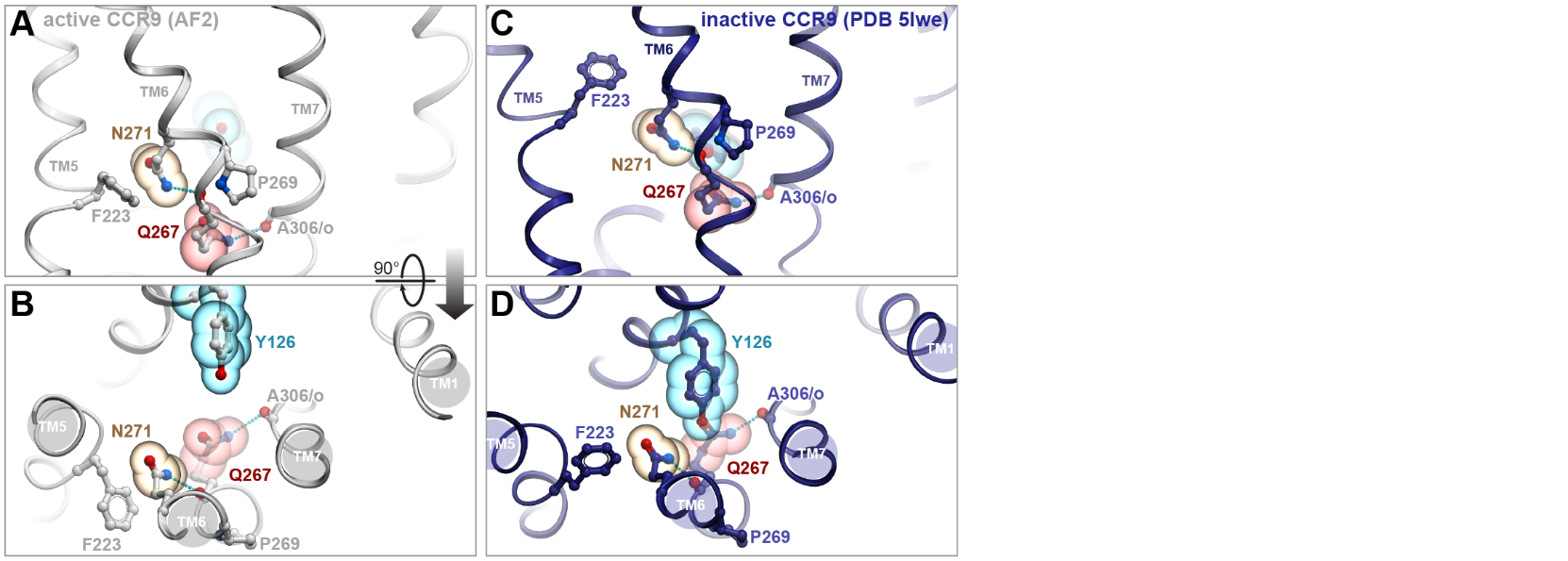

**Figure S15. Pocket floor residues Q267^6.48^ and N271^6.52^ interlock CCR9 TM6 to TM7 and TM5, respectively.** N271^6.52^ pi-stacks with F223^5.47^ of TM5, whereas Q267^6.48^ hydrogen-bonds to the backbone oxygen of A306^7.42^ of TM7. **A and B**, active-state AF2 model. **C and D**, inactive antagonist-bound structure, PDB entry 5LWE. In (**A**) and (**C**), the base of the binding pocket is viewed parallel to the membrane; in (**B**) and (**D**), the same region is viewed perpendicular to membrane from the extracellular side.

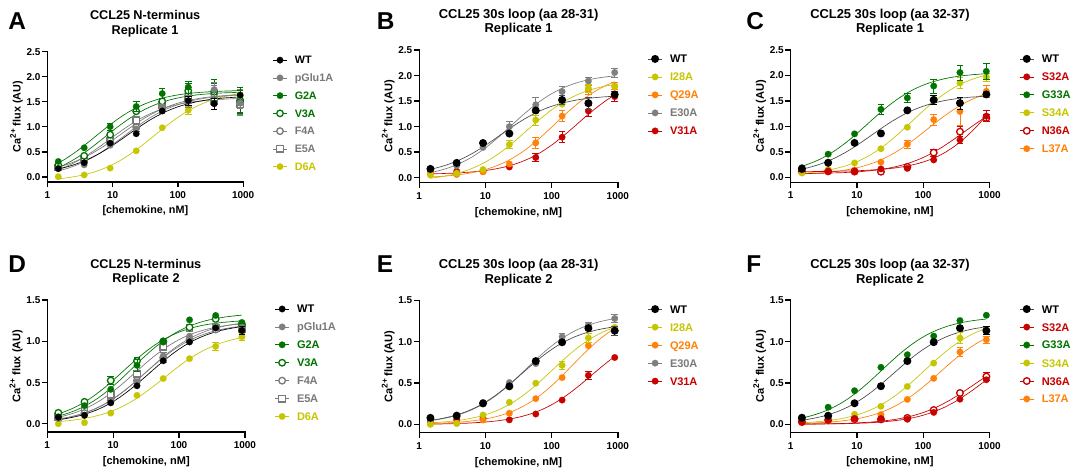

**Figure S16.** **Concentration-dependent responses of MOLT-4 cells to WT CCL25 and the indicated CCL25 mutants in the intracellular Ca^2+^ mobilization assay.** Data points represent mean ± SD for peak height obtained in technical triplicates in 2 independent experiments (experiment 1 (**A-C**) and experiment 2 (**D-F**)).

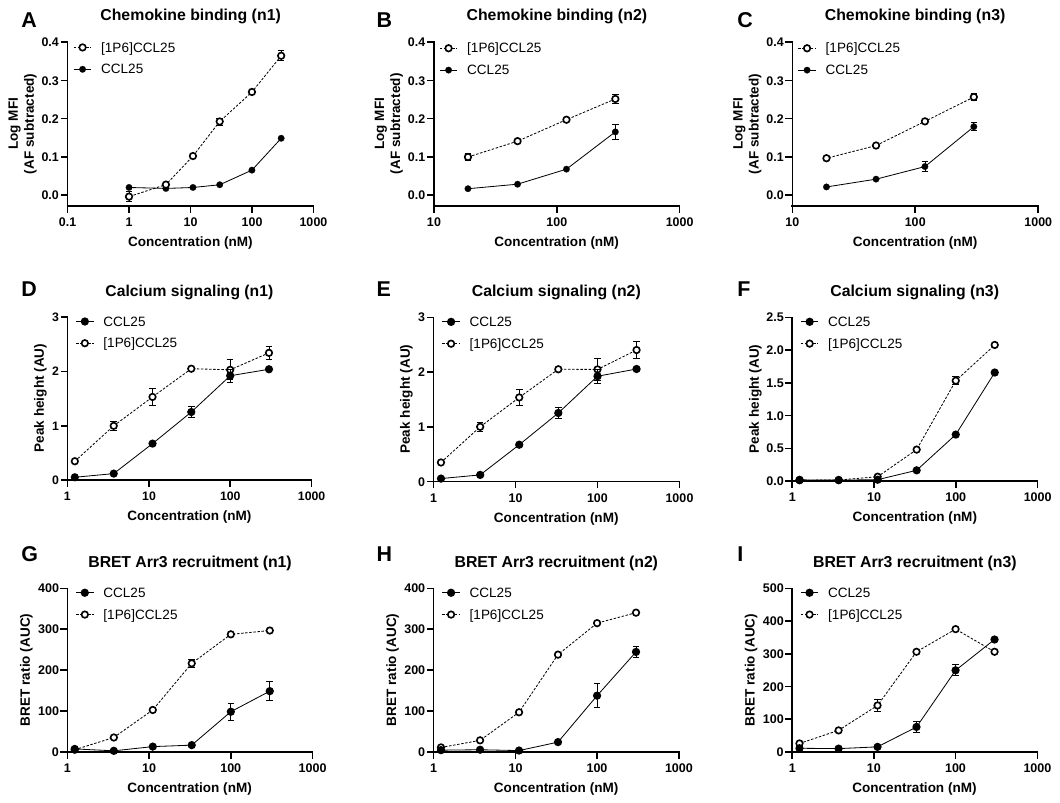

**Figure S17. Concentration-response curves for [1P6]CCL25 binding, Ca^2+^ flux and BRET Arr3 recruitment to WT CCR9 stably expressed in HEK293 cells. A-C.** CCL25 and [1P6]CCL25 binding to CCR9 stably expressed in HEK293 cells. HEK293 cells stably expressing CCR9 were incubated at 4°C for 1h with the indicated concentrations of TAMRA-labelled CCL25 or [1P6]CCL25. Signals are expressed as Log MFI_chemokine_ – Log MFI_AF_. Data represent mean ± SEM of binding signal from triplicate wells. **D-F.** CCL25 and [1P6]-CCL25-induced Ca^2+^ signaling on WT CCR9 expressed in HEK293 cells. Ca^2+^ signals in response to CCL25 and [1P6]CCL25 at the indicated concentrations are shown as mean peak height (**eq. 3**) ± SEM from triplicate wells; the data shown are representative of 3 independent experiments. **G-I.** BRET assays for CCL25 and [1P6]-CCL25-induced Arr3 recruitment on WT CCR9. Data points represent mean ± SEM of BRET signal (**eq. 4**) obtained in triplicate wells.

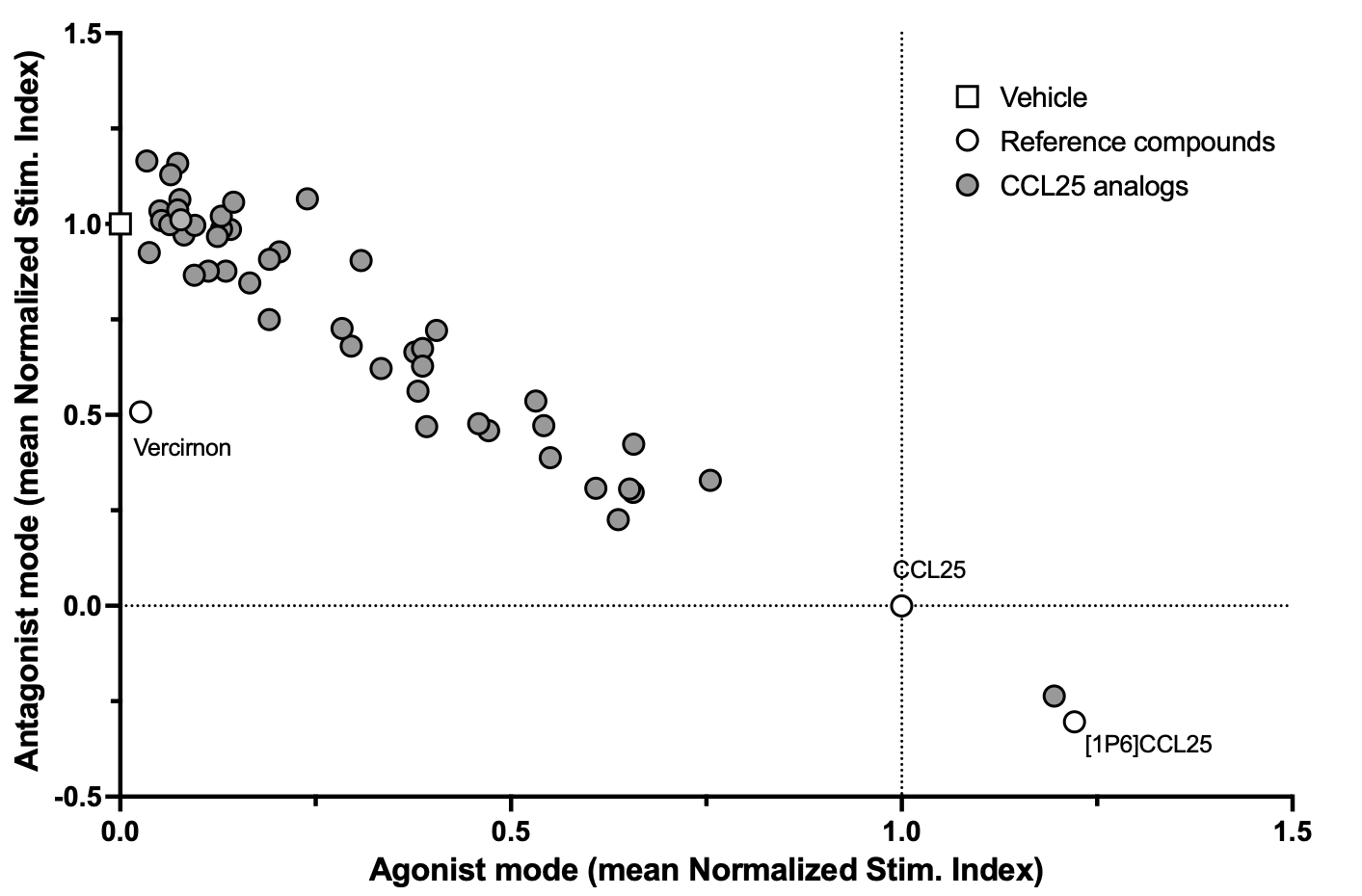

**Figure S18. Activity of the CCL25 N-terminally extended and natural-length analogs discovered by phage-display.** Agonist mode stimulation indexes (SIs) correspond to AUCRC of Ca^2+^ flux responses of MOLT-4 cells to CCL25 analogs. Antagonist mode stimulation indexes (SIs) correspond to AUCRC upon subsequent treatment with CCL25 (100 nM). Vercirnon was used as a positive control antagonist. SIs were normalized by the AUCRC of WT CCL25 (stimulation index set to 1 for agonist mode and to 0 for antagonist mode) and by the AUCRC of vehicle + 100 nM CCL25 mode (SI set to 0 for agonist mode and to 1 for antagonist mode).

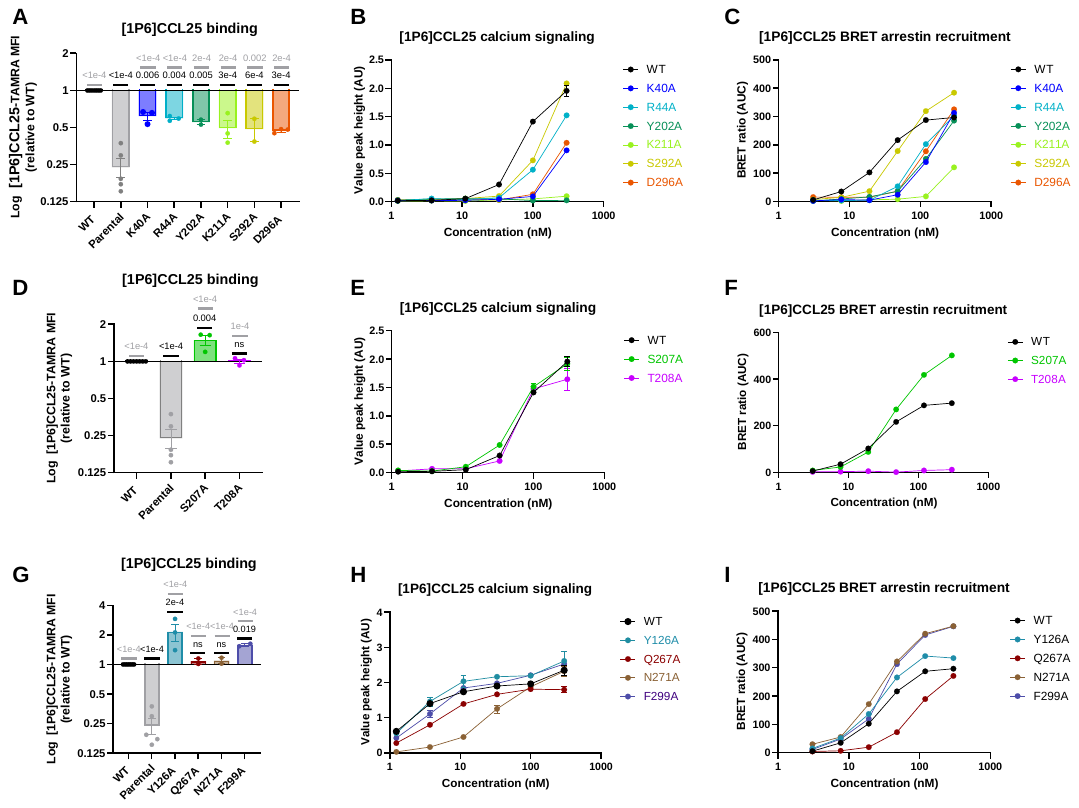

**Figure S19. Binding of [1P6]CCL25 and concentration-response curves for [1P6]CCL25-induced Ca^2+^ flux and BRET Arr3 recruitment to WT and mutant CCR9 stably expressed in HEK293 cells.** Statistical analyses are shown in **Table S6. A, D, G.** [1P6]CCL25 binding to WT CCR9 and CCR9 mutants expressed in HEK293 cells. Cells were incubated at 4°C for 1h with 300 nM of TAMRA-labelled [1P6]CCL25. Bars represent mean ± SEM of the ratio of specific binding signals (**eq. 2**) between the mutant and WT CCR9, measured in 2-3 independent experiments. P-values in comparison to HEK-CCR9 WT and HEK293 parental cells are shown for each mutant in black and grey, respectively. Complete CCL25 binding CRCs are available in **Figs. S20, S21 and S22**. **B, E, H.** [1P6]CCL25-induced Ca^2+^ signaling on WT CCR9 or CCR9 mutants expressed in HEK293 cells. Ca^2+^ signals in response to [1P6]CCL2 at the indicated concentrations are shown as mean peak height (**eq. 3**) ± SEM from triplicate wells; the data shown are representative of 3 independent experiments (complete data set is available in **Figs**. **S23, S24 and S25**). **C, F, I.** BRET assays for [1P6]CCL25-induced Arr3 recruitment on WT CCR9 and CCR9 mutants. Data points represent mean ± SEM of BRET signal (**eq. 4**) obtained in triplicate wells; data shown is representative of 3 independent experiments (complete data set is available in **Figs**. **S23, S24 and S25**).

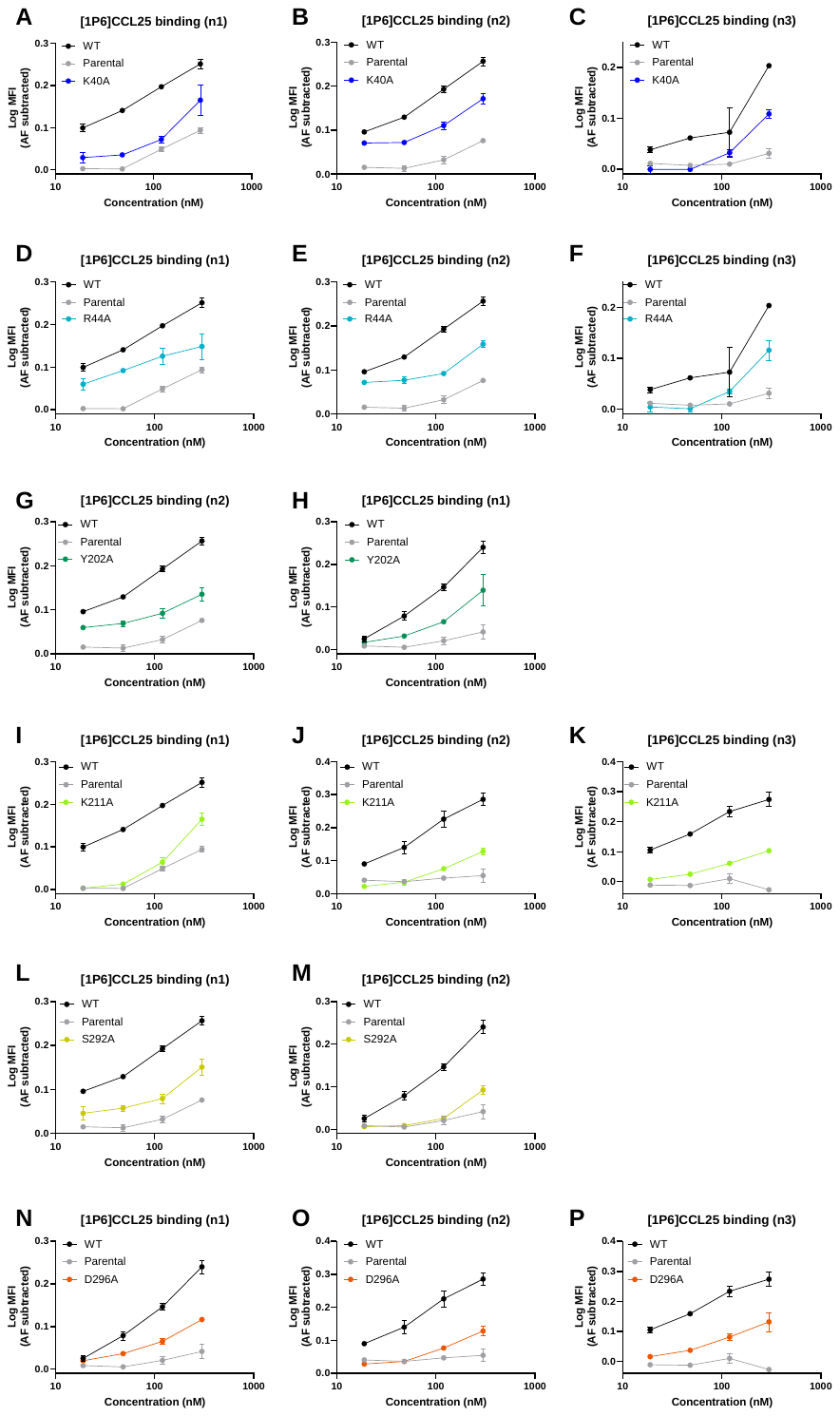

**Figure S20. Concentration-response curves for [1P6]CCL25 binding to K40^1.24^A (A-C), R44^1.28^A (E-F), Y202^ECL2^A (G, H), and K211^5.35^A (I-K), S292^7.28^A (L, M) and D296^7.32^A (N-P) CCR9 mutants compared to CCR9 WT.** Refer to **Fig. S7** legend for details.

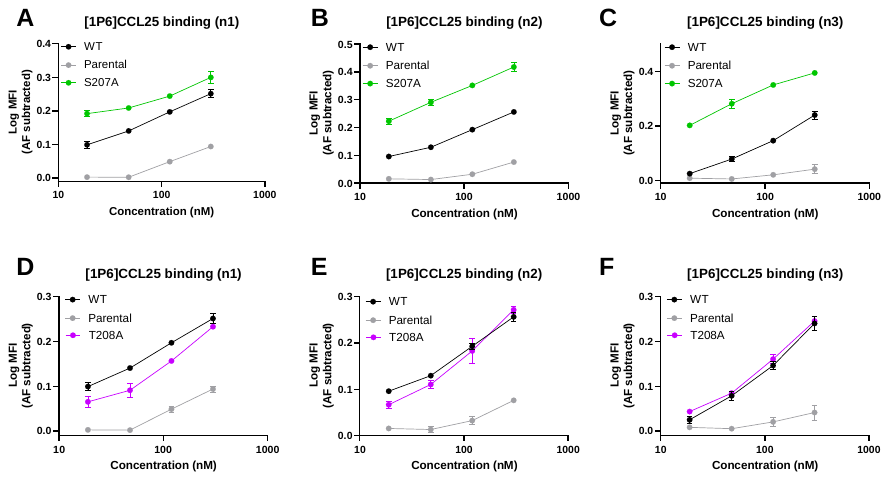

**Figure S21. Concentration-response curves for [1P6]CCL25 binding to S207^5.31^A (A-C) and T208^5.32^A (D-F) CCR9 mutants compared to CCR9 WT.** Refer to **Fig. S7** legend for details.

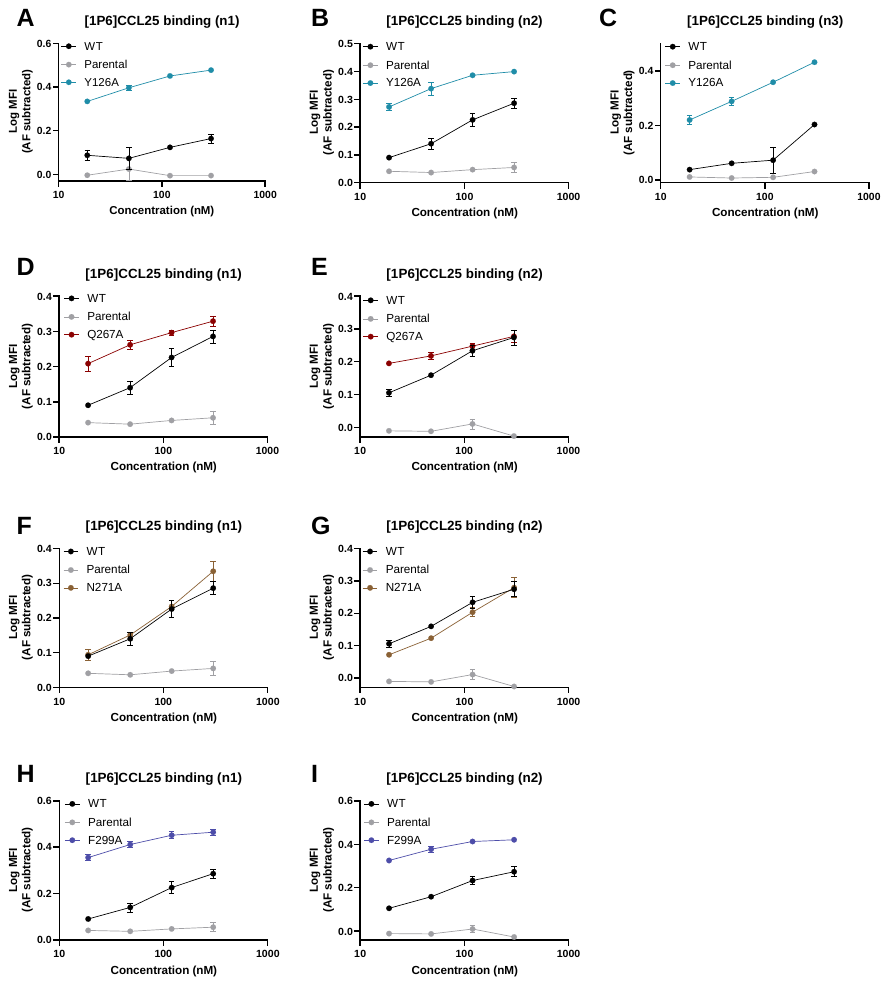

**Figure S22. Concentration-response curves for [1P6]CCL25 binding to Y126^3.32^A (A-C), Q267^6.48^A (D-F), N271^6.52^A (G-I), and F299^7.35^A (J-L) CCR9 mutants compared to CCR9 WT.** Refer to **Fig. S7** legend for details.

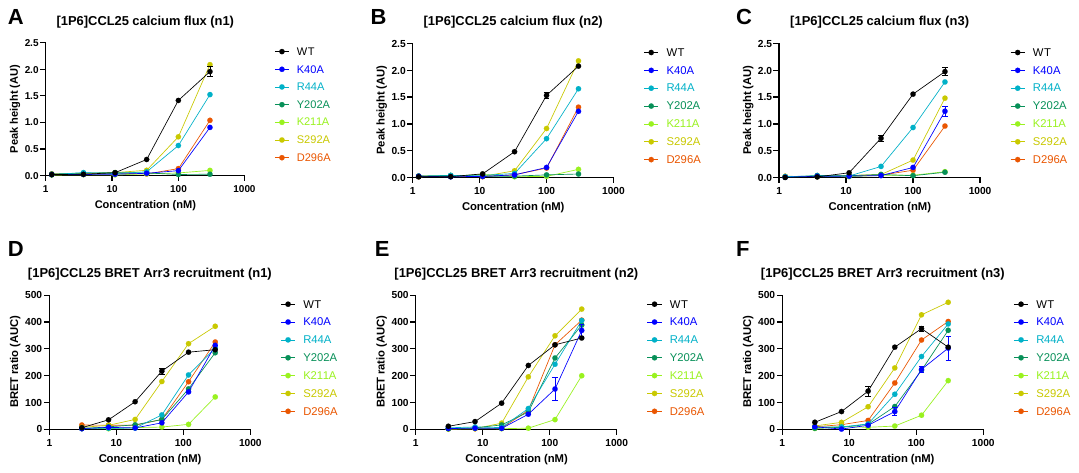

**Figure S23. Concentration-response curves for [1P6]CCL25-induced Ca^2+^ flux (A-C) and BRET Arr3 recruitment (D-F) in K40^1.24^A, R44^1.28^A, Y202^ECL2^A, and K211^5.35^A, S292^7.28^A and D296^7.32^A CCR9 mutants compared to CCR9 WT.** Refer to **Fig. S8** legend for details.

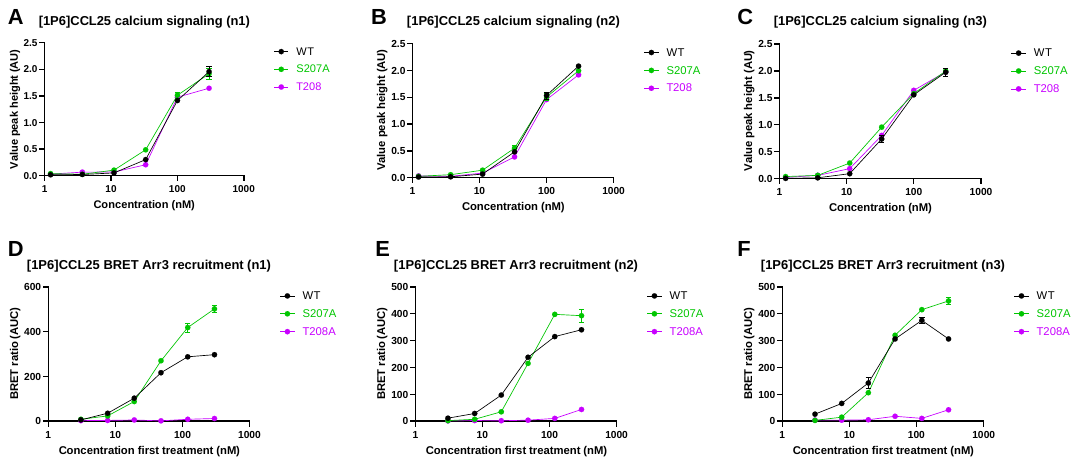

**Figure S24. Concentration-response curves for [1P6]CCL25-induced Ca^2+^ flux (A-C) and BRET Arr3 recruitment (D-F) on S207^5.31^A and T208^5.32^A CCR9 mutants compared to CCR9 WT.** Refer to **Fig. S8** legend for details.

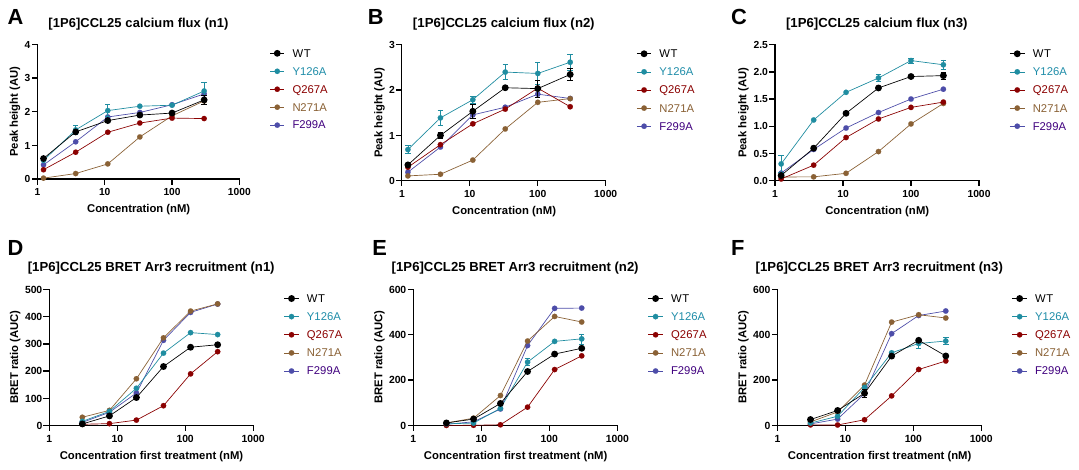

**Figure S25. Concentration-response curves for [1P6]CCL25-induced Ca^2+^ flux (A-C) and BRET Arr3 recruitment (D-F) on Y126^3.32^A, Q267^6.48^A, N271^6.52^A, and F299^7.35^A CCR9 mutants compared to CCR9 WT.** Refer to **Fig. S8** legend for details.

### Supplementary Tables

**Table S1. Key resources and reagents.**

| **Reagent name** | **Manufacturer** | **Catalog number** |
| --- | --- | --- |
| Recombinant CCL25(1-127) | R&D Systems | 9046-TK |
| TAMRA-PEG4-DBCO (Dibenzocyclooctyne-PEG4-Fluor 545) | Sigma-Aldrich | 760773 |
| Amicon Ultra-0.5 Centrifugal Filter 10 kDa MWCO | Millipore | UFC5010 |
| Alexa Fluor 647-conjugated Mouse Anti-Human CD199 (CCR9) mAb | BD Biosciences | 557975; RRID: AB_2073270 |
| HEK293T cells | ATCC | CRL-3216 |
| MOLT-4 cells | ATCC | CRL-1582 |
| DMEM, high glucose, GlutaMAX™ Supplement, pyruvate | Thermo Fisher Scientific | 31966021 |
| RPMI 1640 Medium, GlutaMAX™ Supplement | Thermo Fisher Scientific | 61870036 |
| Fetal bovine serum (FBS) | Thermo Fisher Scientific | A5256701 |
| FluoroBrite™ DMEM | Thermo Fisher Scientific | A1896701 |
| Penicillin-Streptomycin (10,000 U/mL) | Thermo Fisher Scientific | 15140122 |
| JetPRIME® transfection reagent | Polyplus | 101000027 |
| Coelenterazine-h (CTZ-h) | Carbosynth | FC36332 |
| Screen Quest™ Fluo-8 No Wash Ca^2+^ Assay Kit | AAT BioQuest Lubio Science | 36316 |

**Table S2. Constructs and cloning**

| **Construct** | **Plasmid backbone** | **Source lab** | **References** |
| --- | --- | --- | --- |
| Gαi1(91)-RLuc2 | pcDNA3.1(+) | Michel Bouvier, Univ. of Montreal | Quoyer et al. 2013 |
| mVenus-Gβ1 | pcDNA3.1 | Nevin A. Lambert, Univ. of Augusta | Brown et al. 2016; Hollins et al. 2009 |
| Gγ2 | pcDNA3.1 | Aska Inoue, Tohoku Univ. | Inoue et al. 2019 |
| FUGW-CCR9A WT and mutants | FUGW | This work |  |
| FUGW-CCR9A-RLuc8 WT and mutants | FUGW | This work |  |
| FUGW-YFP-arrestin 3 | FUGW | Oliver Hartley, Univ. of Geneva | Martins et al. 2020 |
| psPAX2 | psPAX2 | Patrick Salmon, Univ. of Geneva | Giry-Laterrière et al. 2011 |
| pMD2G | pMD2G | Patrick Salmon, Univ. of Geneva | Giry-Laterrière et al. 2011 |

**Table S3. Statistical analysis of the binding and functional assays on HEK293 cells expressing WT or mutant CCR9 cells and treated with CCL25.**

| **CCL25** | **300 nM CCL25 binding**  **ratio to WT (eq. 1)** | | | **Ca^2+^ flux**  **AUCRC ratio to WT** | | **Arr3 recruitment**  **AUCRC ratio to WT** | |  |
| --- | --- | --- | --- | --- | --- | --- | --- | --- |
| **Condition** | **mean±SEM** | **p-value vs WT CCR9** | **p-value vs parental** | **mean±SEM** | **p-value vs WT** | **mean±SEM** | **p-value vs WT** | **Figs** |
| **WT** | 1.00 ± 0.00 | **n/a** | <0.0001 | 1.00 ± 0.00 | **n/a** | 1.00 ± 0.00 | **n/a** |  |
| **parental** | 0.50 ± 0.05 | <0.0001 | **n/a** | **n/a** | **n/a** | **n/a** | **n/a** |  |
| K40A | 0.84 ± 0.08 | 0.545 | 0.001 | 0.10 ± 0.01 | <0.0001 | 0.12 ± 0.02 | <0.0001 | 2, 3,  S7, S8 |
| R44A | 0.58 ± 0.07 | 0.000 | 0.545 | 0.14 ± 0.02 | <0.0001 | 0.09 ± 0.02 | <0.0001 |  |
| Y202A | 0.81 ± 0.07 | 0.545 | 0.007 | 0.12 ± 0.02 | <0.0001 | 0.19 ± 0.03 | <0.0001 |  |
| K211A | 1.08 ± 0.07 | 0.877 | <0.0001 | 0.07 ± 0.01 | <0.0001 | 0.08 ± 0.04 | <0.0001 |  |
| S292A | 0.66 ± 0.05 | 0.028 | 0.234 | 0.48 ± 0.10 | 0.002 | 0.87 ± 0.10 | 0.508 |  |
| D296A | 0.84 ± 0.03 | 0.545 | 0.001 | 0.14 ± 0.04 | <0.0001 | 0.45 ± 0.04 | 0.006 |  |
| S207A | 1.59 ± 0.15 | 0.004 | <0.0001 | 1.80 ± 0.06 | 0.018 | 1.65 ± 0.22 | 0.124 | 2, 4,  S9, S10 |
| T208A | 0.95 ± 0.04 | 0.877 | <0.0001 | 0.96 ± 0.10 | 0.794 | 0.06 ± 0.01 | <0.0001 |  |
| Y126A | 2.52 ± 0.43 | <0.0001 | <0.0001 | 2.05 ± 0.64 | 0.014 | 2.01 ± 0.26 | 0.021 | 2, 5,  S12, S13 |
| Q267A | 2.06 ± 0.26 | 0.000 | <0.0001 | 1.44 ± 0.38 | 0.325 | 0.29 ± 0.06 | <0.0001 |  |
| N271A | 1.06 ± 0.04 | 0.877 | <0.0001 | 0.18 ± 0.01 | <0.0001 | 2.41 ± 0.44 | 0.005 |  |
| F299A | 2.23 ± 0.24 | <0.0001 | <0.0001 | 1.30 ± 0.29 | 0.467 | 1.52 ± 0.16 | 0.158 |  |

**Table S4. Statistical analysis of the Ca^2+^ flux assay on MOLT-4 cells with WT CCL25 and the indicated CCL25 mutants.** Related to **Fig. 6** and **Fig. S16**.

| **CCL25 ala-scan** | **Ca^2+^ flux** | | | |
| --- | --- | --- | --- | --- |
| **Conditions** | **EC50 (nM)** | **Emax** | **Hill slope** | **p value** |
| **CCL25 WT** | 26.64 | 0.63 | 1.09 | **n/a** |
| CCL25-Z1A | 23.59 | 0.64 | 1.20 | ns |
| CCL25-G2A | 13.04 | 0.68 | 1.27 | 0.017 |
| CCL25-V3A | 12.05 | 0.66 | 1.18 | 0.005 |
| CCL25-F4A | 22.59 | 0.63 | 1.19 | ns |
| CCL25-E5A | 16.65 | 0.62 | 1.35 | ns |
| CCL25-D6A | 46.47 | 0.62 | 1.06 | <0.0015 |
| CCL25-I28A | 61.74 | 0.71 | 0.97 | 0.008 |
| CCL25-Q29A | 123.50 | 0.71 | 1.28 | <0.0015 |
| CCL25-E30A | 30.97 | 0.79 | 0.81 | 0.062 |
| CCL25-V31A | 217.30 | 0.58 | 1.43 | <0.0015 |
| CCL25-S32A | 464.70 | 0.49 | 1.56 | <0.0015 |
| CCL25-G33A | 14.03 | 0.83 | 0.61 | <0.0015 |
| CCL25-S34A | 80.75 | 0.73 | 1.15 | 0.002 |
| CCL25-N36A | 274.70 | 0.43 | 1.62 | <0.0015 |
| CCL25-L37A | 132.10 | 0.66 | 1.10 | <0.0015 |

**Table S5. Activity and sequences of the CCL25 N-terminally extended and natural-length analogs discovered by phage-display.** Z represents pyroglutamate, incorporated during chemical synthesis. Agonist mode stimulation indexes (SIs) correspond to AUCRC of Ca^2+^ flux responses of MOLT-4 cells to CCL25 analogs. Refer to **Fig. S18** legend for details.

|  | | | | **N-terminal sequence** | | | | | | |
| --- | --- | --- | --- | --- | --- | --- | --- | --- | --- | --- |
| **Compound** | **Agonist**  **SI ± SEM** | **Antagonist**  **SI ± SEM** | **№ Replicates** | **0** | **1** | **2** | **3** | **4** | **5** | **6** |
| CCL25 | 1 | 0 | 28 | - | Z | G | V | F | E | D |
| Vercirnon | 0.026 ± 0.003 | 0.508 ± 0.097 | 23 |  | | | | | | |
| CCL25-1P01 | 0.034 ± 0.02 | 1.165 ± 0.08 | 3 | - | Z | G | A | L | R | Q |
| CCL25-1P02 | 0.24 ± 0.038 | 1.066 ± 0.107 | 3 | - | Z | G | V | A | R | N |
| CCL25-1P03 | 0.095 ± 0.042 | 0.866 ± 0.178 | 3 | - | Z | G | V | A | R | R |
| CCL25-1P04 | 0.053 ± 0.018 | 1.009 ± 0.121 | 3 | - | Z | G | V | Q | R | I |
| CCL25-1P05 | 0.078 ± 0.005 | 1.011 ± 0.128 | 3 | - | - | Z | L | G | V | Q |
| CCL25-1P06 | 1.221 ± 0.053 | -0.304 ± 0.094 | 6 | - | Y | Q | A | S | E | D |
| CCL25-1P07 | 0.166 ± 0.029 | 0.846 ± 0.163 | 3 | - | Y | Q | S | R | E | D |
| CCL25-1P08 | 0.135 ± 0.015 | 0.877 ± 0.126 | 3 | - | Y | S | Q | R | E | D |
| CCL25-1P09 | 0.113 ± 0.025 | 0.877 ± 0.12 | 3 | Z | G | A | F | Q | P | D |
| CCL25-1P10 | 0.387 ± 0.025 | 0.628 ± 0.138 | 3 | Z | G | G | F | K | Q | D |
| CCL25-1P11 | 0.191 ± 0.036 | 0.75 ± 0.114 | 3 | Z | G | G | F | Q | W | D |
| CCL25-1P12 | 0.609 ± 0.017 | 0.308 ± 0.108 | 3 | Z | G | F | L | T | A | D |
| CCL25-1P13 | 0.637 ± 0.07 | 0.226 ± 0.059 | 3 | Z | G | G | L | Q | F | D |
| CCL25-1P14 | 0.392 ± 0.035 | 0.469 ± 0.097 | 3 | Z | G | L | L | Q | Q | D |
| CCL25-1P15 | 0.381 ± 0.025 | 0.562 ± 0.139 | 3 | Z | G | Q | L | Q | F | D |
| CCL25-1P16 | 0.295 ± 0.017 | 0.68 ± 0.152 | 3 | K | D | L | Q | F | E | D |
| CCL25-1P17 | 0.471 ± 0.05 | 0.459 ± 0.094 | 3 | L | D | A | Q | F | E | D |
| CCL25-1P18 | 0.656 ± 0.061 | 0.297 ± 0.126 | 3 | T | D | I | Q | F | E | D |
| CCL25-1P19 | 0.55 ± 0.056 | 0.388 ± 0.075 | 3 | V | D | G | Q | F | E | D |
| CCL25-1P20 | 0.652 ± 0.033 | 0.306 ± 0.087 | 3 | V | E | L | Q | F | E | D |
| CCL25-1P21 | 0.334 ± 0.055 | 0.621 ± 0.04 | 3 | E | F | L | R | F | E | D |
| CCL25-1P22 | 0.284 ± 0.018 | 0.726 ± 0.087 | 3 | G | Q | L | K | F | E | D |
| CCL25-1P23 | 0.405 ± 0.01 | 0.722 ± 0.227 | 2 | I | T | Q | R | F | E | D |
| CCL25-1P24 | 0.387 ± 0.009 | 0.674 ± 0.165 | 2 | S | I | Q | R | F | E | D |
| CCL25-1P25 | 0.657 ± 0.068 | 0.423 ± 0.087 | 2 | Z | G | I | Q | F | I | D |
| CCL25-1P26 | 0.542 ± 0.05 | 0.472 ± 0.033 | 2 | Z | G | I | Q | W | I | D |
| CCL25-1P28 | 0.124 ± 0.014 | 0.968 ± 0.102 | 3 | Z | G | I | W | Q | Y | D |
| CCL25-1P29 | 1.195 ± 0.06 | -0.236 ± 0.11 | 2 | Z | G | V | Q | Y | G | D |
| CCL25-1P30 | 0.459 ± 0.084 | 0.477 ± 0.031 | 3 | Z | L | L | W | F | E | D |
| CCL25-1P31 | 0.051 ± 0.006 | 1.034 ± 0.094 | 3 | Z | G | D | I | Q | P | D |
| CCL25-1P32 | 0.037 ± 0.009 | 0.925 ± 0.111 | 3 | Z | G | D | Q | P | I | D |
| CCL25-1P33 | 0.076 ± 0.015 | 1.064 ± 0.135 | 3 | - | R | G | R | Q | E | D |
| CCL25-1P34 | 0.082 ± 0.007 | 0.971 ± 0.07 | 3 | - | R | R | A | E | E | D |
| CCL25-1P35 | 0.095 ± 0.017 | 0.996 ± 0.099 | 3 | - | R | R | K | Q | E | D |
| CCL25-1P36 | 0.145 ± 0.024 | 1.057 ± 0.11 | 2 | - | Z | G | K | S | Q | G |
| CCL25-1P37 | 0.063 ± 0.01 | 0.999 ± 0.121 | 3 | - | Z | G | R | Q | A | Q |
| CCL25-1P38 | 0.074 ± 0.015 | 1.037 ± 0.144 | 3 | - | Z | G | R | S | Q | Q |
| CCL25-1P39 | 0.141 ± 0.006 | 0.985 ± 0.018 | 2 | - | Z | S | K | R | E | D |
| CCL25-1P40 | 0.377 ± 0.044 | 0.665 ± 0.041 | 2 | - | Z | Y | K | Q | E | D |
| CCL25-1P41 | 0.074 ± 0.046 | 1.159 ± 0.139 | 2 | - | Z | G | A | W | W | R |
| CCL25-1P42 | 0.064 ± 0.049 | 1.129 ± 0.147 | 2 | - | Z | G | E | L | H | Q |
| CCL25-1P44 | 0.308 ± 0.005 | 0.904 ± 0.185 | 2 | - | Z | G | Q | W | S | G |
| CCL25-1P45 | 0.129 ± 0.046 | 1.021 ± 0.171 | 2 | Z | G | Q | Y | L | D | D |
| CCL25-1P46 | 0.532 ± 0.042 | 0.537 ± 0.137 | 2 | Z | G | S | Q | L | Q | D |
| CCL25-1P47 | 0.191 ± 0.024 | 0.907 ± 0.06 | 2 | G | R | D | Q | F | E | D |
| CCL25-1P48 | 0.13 ± 0.024 | 0.987 ± 0.169 | 2 | G | R | E | Q | F | E | D |
| CCL25-1P49 | 0.755 ± 0.06 | 0.329 ± 0.208 | 2 | - | V | F | Q | L | E | D |
| CCL25-1P50 | 0.204 ± 0.044 | 0.927 ± 0.145 | 2 | - | V | Q | R | L | E | D |

**Table S6. Statistical analysis of the binding and functional assays on HEK293 cells expressing WT or mutant CCR9 cells and treated with [1P6]CCL25.** Related to **Fig. 8** and **Figs. S19** to **S25.**

| **[1P6]**  **CCL25** | **300 nM [1P6] CCL25 binding**  **ratio to WT (eq. 1)** | | | **Ca^2+^ flux**  **AUCRC ratio to WT** | | **Arr3 recruitment**  **AUCRC ratio to WT** | |
| --- | --- | --- | --- | --- | --- | --- | --- |
| **Condition** | **mean±SEM** | **p-value vs WT CCR9** | **p-value vs parental** | **mean±SEM** | **p-value vs WT** | **mean±SEM** | **p-value vs WT** |
| **WT** | 1.00 ± 0.00 | **n/a** | <0.0001 | 1.00 ± 0.00 | **n/a** | 1.00 ± 0.00 | **n/a** |
| **parental** | 0.24 ± 0.04 | <0.0001 | **n/a** | n/a ± n/a | **n/a** | n/a ± n/a | **n/a** |
| K40A | 0.62 ± 0.04 | 0.006 | <0.0001 | 0.26 ± 0.02 | <0.0001 | 0.44 ± 0.01 | <0.0001 |
| R44A | 0.59 ± 0.01 | 0.004 | <0.0001 | 0.57 ± 0.03 | <0.0001 | 0.59 ± 0.03 | 0.0004 |
| Y202A | 0.55 ± 0.03 | 0.005 | 0.0048 | 0.06 ± 0.00 | <0.0001 | 0.53 ± 0.06 | <0.0001 |
| K211A | 0.49 ± 0.08 | 0.000 | 0.0003 | 0.06 ± 0.00 | <0.0001 | 0.15 ± 0.01 | <0.0001 |
| S292A | 0.49 ± 0.10 | 0.001 | 0.002 | 0.59 ± 0.12 | <0.0001 | 0.94 ± 0.01 | 0.727 |
| D296A | 0.47 ± 0.01 | 0.000 | 0.0003 | 0.26 ± 0.02 | <0.0001 | 0.65 ± 0.06 | 0.003 |
| S207A | 1.49 ± 0.15 | 0.064 | <0.0001 | 1.10 ± 0.03 | 0.597 | 1.12 ± 0.11 | 0.727 |
| T208A | 1.00 ± 0.04 | 0.795 | <0.0001 | 0.99 ± 0.05 | 0.920 | 0.04 ± 0.01 | <0.0001 |
| Y126A | 2.15 ± 0.44 | 0.000 | <0.0001 | 1.19 ± 0.04 | 0.383 | 1.11 ± 0.06 | 0.727 |
| Q267A | 1.08 ± 0.07 | 0.767 | <0.0001 | 0.76 ± 0.05 | 0.065 | 0.54 ± 0.01 | <0.0001 |
| N271A | 1.09 ± 0.08 | 0.767 | <0.0001 | 0.51 ± 0.06 | <0.0001 | 1.45 ± 0.05 | 0.013 |
| F299A | 1.58 ± 0.04 | 0.019 | <0.0001 | 0.89 ± 0.06 | 0.571 | 1.36 ± 0.06 | 0.038 |
